# Gaze reinstatement during working memory for natural scenes

**DOI:** 10.1101/2025.09.29.678115

**Authors:** Yueying Dong, Yun-chen Hung, Lana Gaspariani, Anastasia Kiyonaga

## Abstract

The information we hold in mind with working memory (WM) may propagate beyond the cortex and out to the peripheral motor system. For instance, directionally biased hand and eye movements can express simple visuo-spatial features during WM maintenance. If such sensorimotor inflections are a form of WM representation, they may also express more multifaceted and task-adaptive information than just spatial location. Here, we ask whether gaze patterns carry item-specific detail about complex stimuli held in mind, and whether gaze prospectively adapts to the behavioral task demands. To do this, we tracked human eye movements (male and female) during WM for naturalistic images, and we manipulated which image features were most task-relevant (visual or semantic). In two experiments, we found that the eyes moved in complex, item-specific spatiotemporal sequences during WM maintenance. Delay period gaze patterns retraced the scanpaths from stimulus encoding, echoing the ‘gaze reinstatement’ that aligns with neural pattern reactivation during long-term memory retrieval. Therefore, gaze patterns during WM may reinstate visual encoding processes for maintenance. We also used time-resolved representational similarity analyses, guided by convolutional neural network modeling, to test the specificity of WM gaze patterns across a trial. We found that gaze representations were more distinct when the task prioritized visual detail, and also ramped up in anticipation of the WM probe. Therefore, oculomotor WM signals are malleable to when and how the content will be used. These results suggest that the earliest levels of visual processing reflect prospectively-oriented and functionally flexible WM content representations.

**Significance:** The eyes are an outwardly accessible extension of the brain. They are the first stop for visual processing, but may also offer a rich window into underlying cognition. For instance, pupil dilation can reflect changing cognitive load, and eye movement biases can reflect basic spatial information about content held in mind with working memory. Working memory is a critical neuro-cognitive function, but its physiological bases remain under debate. Here, we show that eye movements code for more than simple spatial WM, and instead express intricate patterns that read out the complex, real-world stimuli held in mind. This work shows how cognitive states influence oculomotor functions, and also advances theories of WM representation to include the earliest stages of sensory processing.

## Introduction

Flexible goal-directed cognition relies on the ability to guide behavior based on task-relevant information that is held ‘in mind.’ Decades of research have sought to unveil the hidden mental representations that support this working memory (WM) function (Christophel et al., 2017; Leavitt et al., 2017; Sreenivasan et al., 2014). Prominent WM theories assert that feature-specific WM content is stored in sensorimotor cortical activations (D’Esposito & Postle, 2015; Serences, 2016), but newer evidence suggests that sensorimotor signals outside the cortex can also carry WM information. For instance, small gaze biases veer toward locations in memorized visual space, and pupil size tracks the remembered brightness of a visual WM stimulus (van Ede et al., 2019; Zokaei et al., 2019). Such ocular WM signatures may provide a valuable window into internal goal states, and into the linkage between WM and motor systems. However, the specific information contained in oculomotor inflections, and their functional sensitivity to task demands, remain unclear.

A growing body of work has begun to shed light on the WM-related information that is expressed in eye movements. Micro-saccades are found to be reliably biased toward the spatial location where a task-relevant visual WM stimulus was encoded (Liu et al., 2022; van Ede et al., 2019). Gaze and saccade biases can also track imagined geometric shapes (Laeng et al., 2014), and the remembered angle of oriented objects (Linde-Domingo & Spitzer, 2024). These tasks may all be considered fundamentally spatial, however, leaving it unclear as to whether oculomotor WM signatures carry more complex feature information than mere spatial tagging.

Long-term memory (LTM) research provides a clue as to the complexity the eyes might convey during WM. Namely, during LTM retrieval, eye movements can retrace the spatiotemporal sequences that occurred during stimulus encoding (Johansson & Johansson, 2014; Wynn et al., 2019). This phenomenon is termed ‘gaze reinstatement’, and it may functionally relate to retrieval success (Johansson & Johansson, 2014; Wynn et al., 2020). Gaze patterns follow similar principles to neural reinstatement and have been linked to hippocampal retrieval processes (Nau et al., 2025; Kragel & Voss, 2022). If WM operates over similar visual representations as LTM (Cowan, 1999; Fukuda & Woodman, 2017; Schurgin, 2018), eye movements during WM may also represent rich visual detail in their spatio-temporal patterns.

However, the detail represented in gaze patterns may also depend on the task at hand. WM is tightly intertwined with action (Boettcher et al., 2021; Miller et al., 2020; Olivers & Roelfsema, 2020), and the same information may be maintained with different neural activation patterns under different expected task demands (Nobre & Stokes, 2019; Lee et al., 2013; Bae & Chen, 2024; Henderson et al., 2022). While the quality of neural WM representations is typically gauged by the correspondence between encoded stimuli and decoded features (Ester et al., 2013), many real-world tasks may be accomplished with coarser WM codes. If WM is geared to guide near future behavior, the WM representation should match the resolution required by the task. For instance, we might expect a richer range of eye motion during WM for visual detail vs. a categorical label.

Here, we address two main questions about oculomotor WM signatures. First, what information do they convey? Specifically, we ask whether delay period eye movements retrace the complex, item-specific patterns made during encoding. Second, are they flexible to prospective task demands? We ask whether WM gaze patterns retrospectively follow the physical properties of encoded stimuli or prospectively adapt to the task context. To do so, we manipulate whether visual detail or semantic stimulus properties are emphasized at test, and whether that test condition is known before or after stimulus encoding. We evaluate the alignment between gaze sequences from encoding to WM maintenance, and whether they vary when different maintenance strategies are emphasized. The findings reveal fine specificity in the inner mental content that is expressed in the eyes, and a sensitive coupling between WM maintenance and motor systems.

## Materials and Methods

### Overview

Across two experiments, we manipulated how WM content would be used for a subsequent match-to-sample task. In both experiments, participants were asked to remember a naturalistic image over a delay of several seconds, and they were cued as to whether the visual or semantic image properties would be most task relevant (**Figure 1A**). On each trial, a color cue indicated whether the WM probe would test on either the visual detail or the category of the sample image. In **Experiment 1**, the color cue appeared before the WM sample stimulus (**precue** task, n = 41), whereas in **Experiment 2** the cue appeared after the WM sample encoding period (**retrocue** task, n = 41). We tracked eye movements at 1000Hz during both experiments, to test whether spatiotemporal gaze patterns convey image-specific information about WM content, and whether they are flexible to the WM task demands.

**Figure 1.**
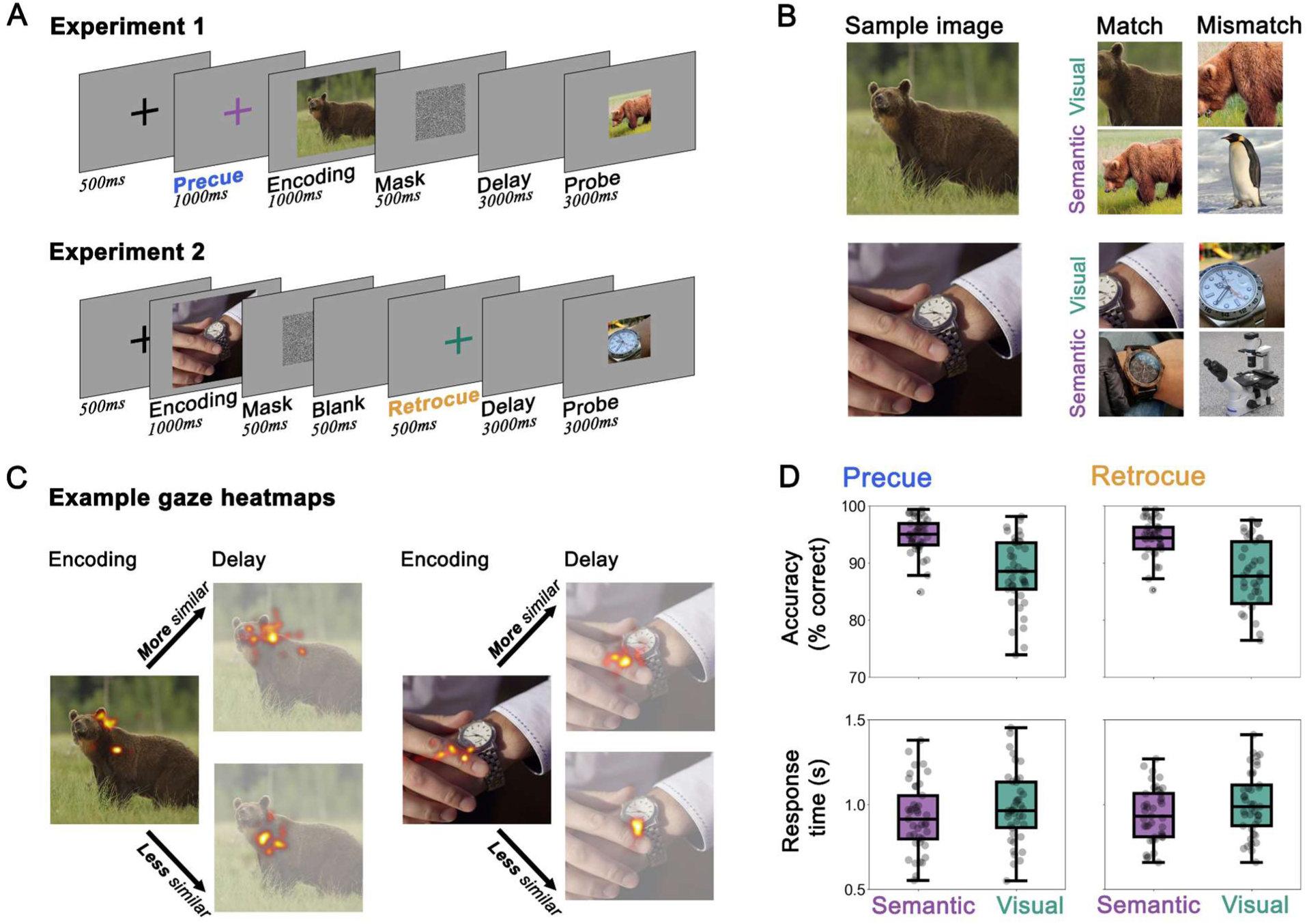
Experiment design and memory performance for both experiments. (**A**) Task schematics for Experiment 1 (precue) and Experiment 2 (retrocue). In both experiments, a color cue indicated whether the probe would test for the visual detail (green) or semantic category (purple) of the WM sample image. However, in the precue task the cue appeared before encoding, whereas in the retrocue task it appeared after encoding. (**B**) Example WM sample stimuli and memory probes under the different trial conditions. (**C**) Example gaze heatmaps from encoding and delay epochs during WM for select images. The delay epoch examples show trials where the gaze heatmaps would be considered either more similar (top) or less similar (bottom) to the gaze during encoding of the same image. Yellower shades represent a longer dwell time. (**D**) Working memory recognition performance for both experiments and trial conditions. Accuracy reflects % correct and response time reflects the duration between probe onset and button press. Data points reflect individual subject means, the box reflects the interquartile range, and the horizontal line reflects the group median.

We first used sequence-alignment algorithms to test whether delay-period gaze patterns matched the patterns from stimulus encoding. Then, to capture the temporal evolution of stimulus information in gaze patterns under various conditions, we also used time-resolved representational similarity analyses. We used a deep convolutional neural network to evaluate the visual similarity between our WM sample images, and we compared that model-derived stimulus similarity with our empirically-derived gaze similarity, as it evolved across a trial.

### Participants

The experiments were approved by the UC San Diego Office of IRB Administration (OIA). Participants were adults (over 18 years of age) with normal or corrected to normal (via contact lens) vision, recruited via UCSD’s SONA platform. Participants provided informed consent, and received course credit as compensation. A session took 60 - 80 minutes to complete.

We recruited 42 participants for Experiment 1 (precue), and 41 for Experiment 2 (retrocue). We based our sample size on related studies that find reliable effects when testing similar multivariate analyses of eye movement data (Linde-Domingo & Spitzer, 2024; Yang et al, 2025). One participant was excluded from the precue experiment for data quality reasons (see *Preprocessing* for Eyetracking QA details), and no participants were excluded from the retrocue experiment. In total, 41 participants were included in precue experiment analyses (*27 females, 12 males, 1 declined to report, 1 other, mean age = 22.84 ± 3.85*) and 41 in retrocue experiment analyses (*34 females, 5 males, 1 declined to report, 1 other, mean age = 20.31 ± 2.55*).

### Stimuli and task procedure

The experiments were programmed and presented in Psychopy, v2023.2.3. Participants completed the task in a dimly lit room on a 1920 x 1080 monitor (24 inch, 60.96cm) with 60hz refresh rate. Participants sat 60cm away from the screen center, their heads stabilized by a chinrest.

WM sample stimuli were 40 unique naturalistic images from the *THINGS* database (Hebart & Stoinski, 2019).Two images were taken from each of 20 randomly-selected categories (e.g., bears, watches, etc). The image set was the same for all participants, to obtain comparable gaze measures across participants.

In both experiments, each trial started with a 500ms fixation, where a black cross appeared at the center of the screen (128 x 128 pixels, subtending 3.5° visual angle). In the **precue experiment** (**Figure 1A**, top panel), the color of the cross then changed to either green or purple. The color of the cross cued which stimulus feature would be tested at the end of the trial. This precue remained onscreen for 1000ms. If the precue (the cross) was green, the WM probe would test for visual detail of the sample image, but if the precue was purple, the probe would test for the semantic category of the sample stimulus. The color cues were designed to be equiluminant to each other and to the neutral grey background (green: RGB 0, 128, 0; purple: RGB 128, 0, 128). During practice, participants were instructed and trained to learn the meaning of the cues.

Following the precue, a WM sample image (540 x 540 pixels; subtending 14° visual angle) was centrally displayed for 1000ms. After the sample image, a noise mask appeared for 500ms, and was then followed by a 3000ms blank memory delay. Lastly, at the end of each trial, a probe image appeared for 3000ms, and participants judged whether it was a match or non-match to the sample image. Probe images subtended 7° visual angle (270 x 270 pixels). On *visual* trials (i.e., green cue), the probe was a cropped segment of either the sample image (match) or a different image from the same category (non-match). For example, when remembering an image of a bear, a match would be cropped from the exact same bear image (**Figure 1B**). The crop was sized at 20% of the original image and was selected from a randomly-drawn location within a circle around the image center. The segment was then scaled up to 270 x 270 for visibility. Showing only a segment of the full image was designed to encourage a more visual maintenance strategy for fine image details.

On *semantic* trials (i.e., purple cue), the probe image was never an exact match to the sample image, but was drawn from either the same category as the sample (match) or an image from a different category altogether (non-match). For example, when remembering an image of a bear, a match would be an image of a different bear (**Figure 1B**). The *semantic* probes were uncropped, but were scaled down to 270 x 270 pixels to match the size of the *visual* probes. Non-match probe stimuli were drawn from outside the sample stimulus set. In both conditions, participants made a keyboard button press to indicate whether the probe was a match or non-match to the sample. Match and non-match probes, as well as *visual* vs. *semantic* trials, occurred equally often and in random order. Thus, to successfully perform the task required either maintenance of precise visual detail or the broader semantic category.

The trial structure of the retrocue experiment (**Figure 1A**, bottom panel) broadly followed that of the precue experiment, except for one critical difference: instead of being told before encoding what stimulus feature would be tested, participants were retroactively cued after the sample stimulus had been encoded. Each trial started with a 500ms fixation, followed by a 1000ms sample encoding period. Next the mask was shown for 500ms, then a 500ms blank screen. Following the blank screen, the retrocue appeared as a color cross. As in the precue experiment, the color (purple or green) indicated the task relevant feature (*visual* or *semantic*). After a 3000ms memory delay, participants completed the same probe task as in the precue experiment.

Both experiments consisted of 320 trials, divided into 16 blocks of 20 trials each. Each unique image was presented 8 times in total – 4 times in each the *visual* and *semantic* conditions. Within that constraint, images were drawn from the stimulus set in random order for each trial. One complete session of either experiment contained 160 *visual* and 160 *semantic* trials.

### Eye-tracking analysis

We implemented continuous eye tracking throughout both experiments via a desk-mounted Eyelink Portable Duo, recording at 1000hz (SR Research Ltd.). We performed a 13-point calibration using the built-in calibration module, and performed a drift check at the beginning of every block (every 20 trials) to ensure that central fixation was maintained. Data was exported using the EyeLink Data Viewer software package (Version 4.3.1).

#### Preprocessing

The eye-tracking preprocessing procedure was similar to previous work, where blink artifacts were detected as outliers in the velocity space of pupillary change (Dong & Kiyonaga, 2024). The data were first transformed into *Trial x Time* format, and for each trial, we obtained a smoothed derivative array, 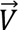, from the original pupil size, 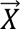

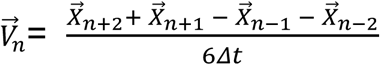

Next, we defined blinks as any time point that exceeded a velocity threshold

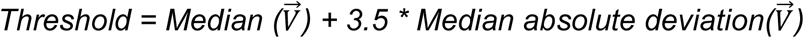

The timepoints exceeding this threshold were marked as blinks, and each was padded with 50ms on either side. This produced a blink mask in the gaze data, and we removed the data during these periods. The missing gaze data were interpolated using linear, nearest-neighbor, or no interpolation if the gap was too large (Dong & Kiyonaga, 2024). After interpolation, a trial was discarded if it still had more than 20% data missing from the main epoch of interest – the 5.5s from fixation offset to the end of the memory delay. Participants were excluded if they had more than 40% trials discarded, either due to poor eye-tracking quality or missing probe responses. After excluding one subject for poor data quality, trial-wise data cleaning retained 91.3% of data in the precue experiment (203 - 320 trials per participant), and 93.5% of data in the retrocue experiment (255 - 319 trials per participant).

#### Saccade detection

Similar to detecting blinks from the pupil array, saccadic activity was detected from the gaze array as outliers in velocity space. We adopted the approach outlined in Engbert & Kliegl (2003). To reduce aliasing for saccade detection, we smoothed the gaze data using a rolling window average of size 11ms, then calculated the gaze velocity on the horizontal and vertical axis 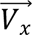, 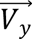. The 2d velocity was then calculated as

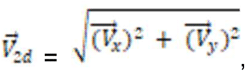

Saccades were outliers on this 2d velocity space that exceeded the threshold of median 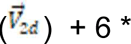 median absolute deviation 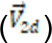. Additionally, we set a minimum shift distance of 8 pixels for a movement to be considered saccadic, equating to 0.4% change relative to the screen size and 0.21° visual angle. These thresholds followed those determined by prior work (Dong & Kiyonaga, 2024). We also calculated the Working memory gaze reinstatement frequency of saccades, for a control analysis (**Figure 2E**), by applying a rolling window filter. Here, FN represents the saccade frequency at Nth window.

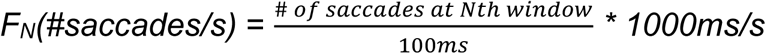

**Figure 2.**
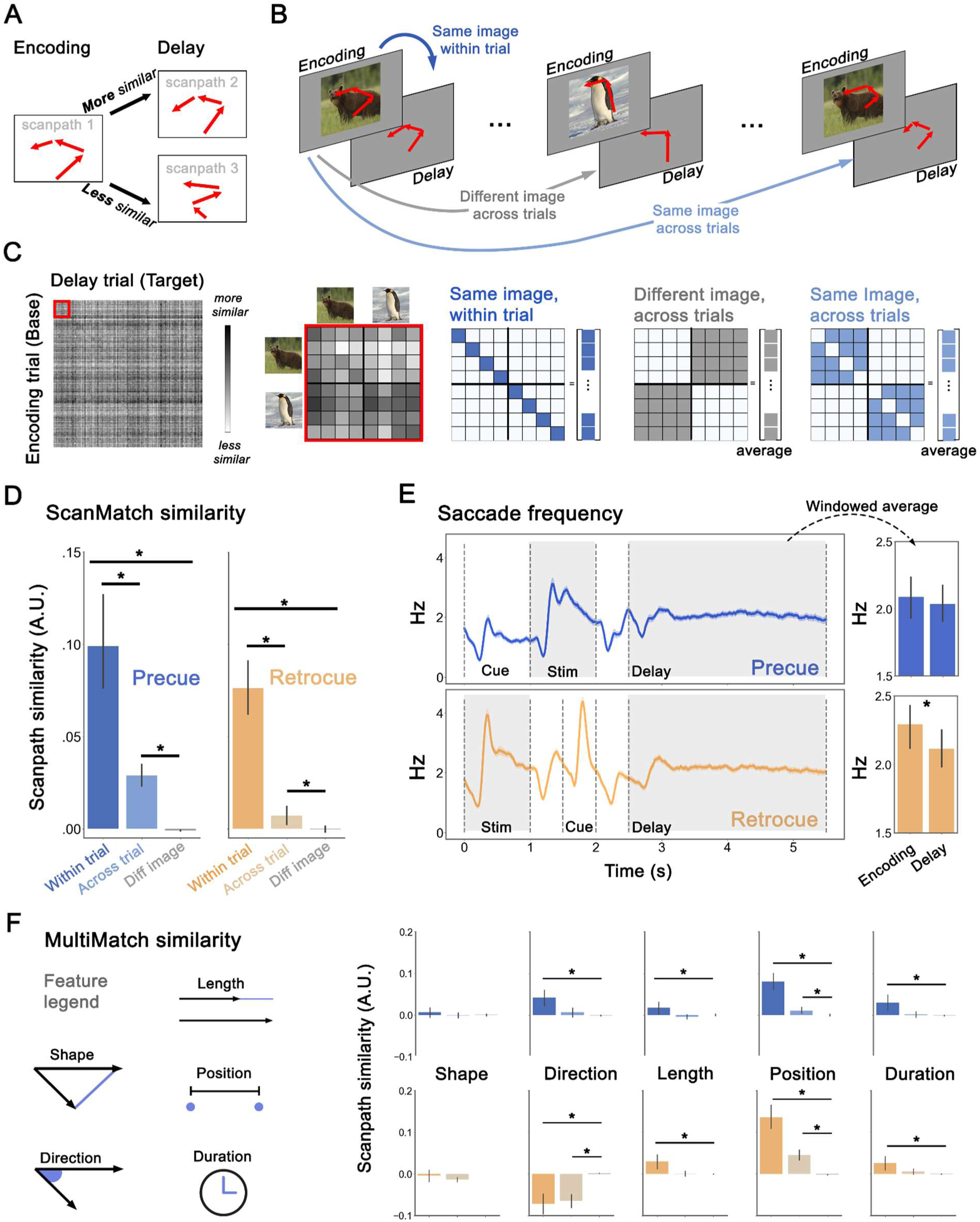
Scanpath method and results for both experiments. (**A**) A schematic illustration of similar vs. dissimilar scanpaths from encoding to delay. (**B**) Schematics illustrating the three types of encoding-delay scanpath similarity that were compared: within the same trial (“within trial”), across different trials where different images were remembered (“different image”), and across different trials where the same image was remembered (“across trial”). (**C**) Left: The subject-averaged scanpath similarity matrix (ScanMatch algorithm) from 320 trials in the precue experiment. Diagonal values represent within-trial comparisons between encoding and delay scanpaths. Darker shades represent more similarity between encoding-delay scanpath pairs. Middle: A subset of the similarity matrix for two images, in just the visual condition. Each image appeared 4 times in each of the visual and semantic trial conditions (8 times total per experiment). Right: Masks applied to the scanpath similarity matrix to extract the three scanpath comparison conditions (“within trial”, “different image”, and “across trial”). Shaded regions represent 1, and white space represents 0. (**D**) The mean of individual subject scanpath similarity scores (ScanMatch algorithm) across comparison conditions and experiments. Note the similarity matrices were normalized within subject to account for baseline variability. Horizontal lines across the top of the plot indicate a main effect of comparison condition, whereas shorter lines accompany significant post-hoc pairwise comparisons. (**E**) Left: Average saccade frequency, over the time course of the trial, in both experiments. Right: Windowed average of saccade frequency for the encoding and delay epochs, in both experiments (areas shaded in grey on the frequency timecourse). (**F**) Scanpath similarity scores for five geometric dimensions (MultiMatch algorithm), across comparison conditions. Shape and direction measure the vector distance and angular distance between aligned saccade vectors, length measures saccade amplitudes, position measures the Euclidean distance between aligned fixations of the saccade vectors, and duration measures the fixation dwell time. Across all panels, error bars reflect 95% CI, and shaded error bands reflect ± 1SEM.

#### Scanpath analysis

A scanpath is a series of fixations. To transform the continuous gaze array into scanpaths, we first extracted the identified saccades for the two epochs of interest: the stimulus encoding and memory delay periods. We then extracted fixation locations (the average eye position on the horizontal and vertical axes between each detected saccade), as well as the fixation duration (dwell time in between each saccade (Coutrot et al., 2018; Cristino et al., 2010; Jarodzka et al., 2010) within each epoch. We did this for every trial, resulting in an encoding and a delay scanpath for each. These scanpaths were then compared using the analysis approaches described below.

Two prevalent approaches to scanpath analysis can be loosely classified as string-based and vector-based (Coutrot et al., 2018; Fahimi & Bruce, 2021).The string-based method represents scanpaths as sequences of letters, each corresponding to a region on the monitor screen. Similarity is then quantified between two strings (Cristino et al., 2010; Fahimi & Bruce, 2021), yielding a single similarity score for each comparison. The vector-based method measures the similarity across multiple vector geometry features, yielding multiple distinct scores (Dewhurst et al., 2012). Both approaches assess the spatio-temporal information in eye-movement patterns, but they do so using different underlying metrics.

For the string-based approach, we used a MATLAB implementation of the ScanMatch algorithm (Cristino et al., 2010). The ScanMatch similarity is the normalized output from a sequence alignment algorithm (Needleman & Wunsch, 1970). This algorithm takes two scanpaths as input and produces a single similarity value representing the degree of similarity between them. We compared scanpaths for every encoding-delay pair. In other words, every trial’s encoding period was compared to the delay period of the same trial, as well as the delay periods from every other trial. This generated a 320 x 320 matrix for every participant, with encoding trials as rows and delay trials as columns (**Figure 2C**). We then z-score normalized each similarity matrix to remove across-participant baseline variability. Diagonal values index the scanpath similarity from encoding to delay in the same trial trial, which we used to measure the strength of gaze reinstatement during the WM delay within a trial (**Figure 2C**, “same image, within trial”). Off-diagonal values for the same image indicated the extent to which image-specific eye-movement patterns during the delay generalized across trials (**Figure 2C**, “same image, across trials”). Off-diagonal values for different images served as a baseline comparison reflecting whether image-general encoding-delay gaze patterns were shared across trials, regardless of the specific stimulus (**Figure 2C**, “different image, across trials”). Since each unique image was presented multiple times in the experiment, scores were averaged across columns for the “different image” and “same image, across trials” comparisons. This yielded three encoding-delay similarity scores for every trial: “within trial”, “across trial”, and “different image”.

Supplementing the string-based analyses (**ScanMatch**), we also used a Python implementation of the vector-based **MultiMatch** algorithm, multimatch-gaze (Dewhurst et al., 2012; Jarodzka et al., 2010; Wagner et al., 2019). Here, instead of letter sequences, scanpaths were represented as vectors and evaluated along five dimensions: shape, length, position, direction, and duration. We completed the same encoding-delay comparisons with both the string-based **ScanMatch** and vector-based **MultiMatch** scores. While the former yielded just one similarity score for each comparison, the latter output five scores for the five dimensions.

To evaluate whether scanpath similarity changed in response to visual and semantic task demands, we then sorted the similarity matrix by whether the encoding scanpath (that was shared by all encoding-delay pairs in a row) was from a *visual* or *semantic* trial. We then compared those encoding epochs only to delay epochs of the same trial condition (**Figure 3A**). The three scanpath comparisons (“within trial”, “across trial”, and “different image”) were thus partitioned into *visual* or *semantic* trial categories. We applied the same procedure to the ScanMatch and Multimatch matrices (**Figure 3B** and **3C**).

**Figure 3.**
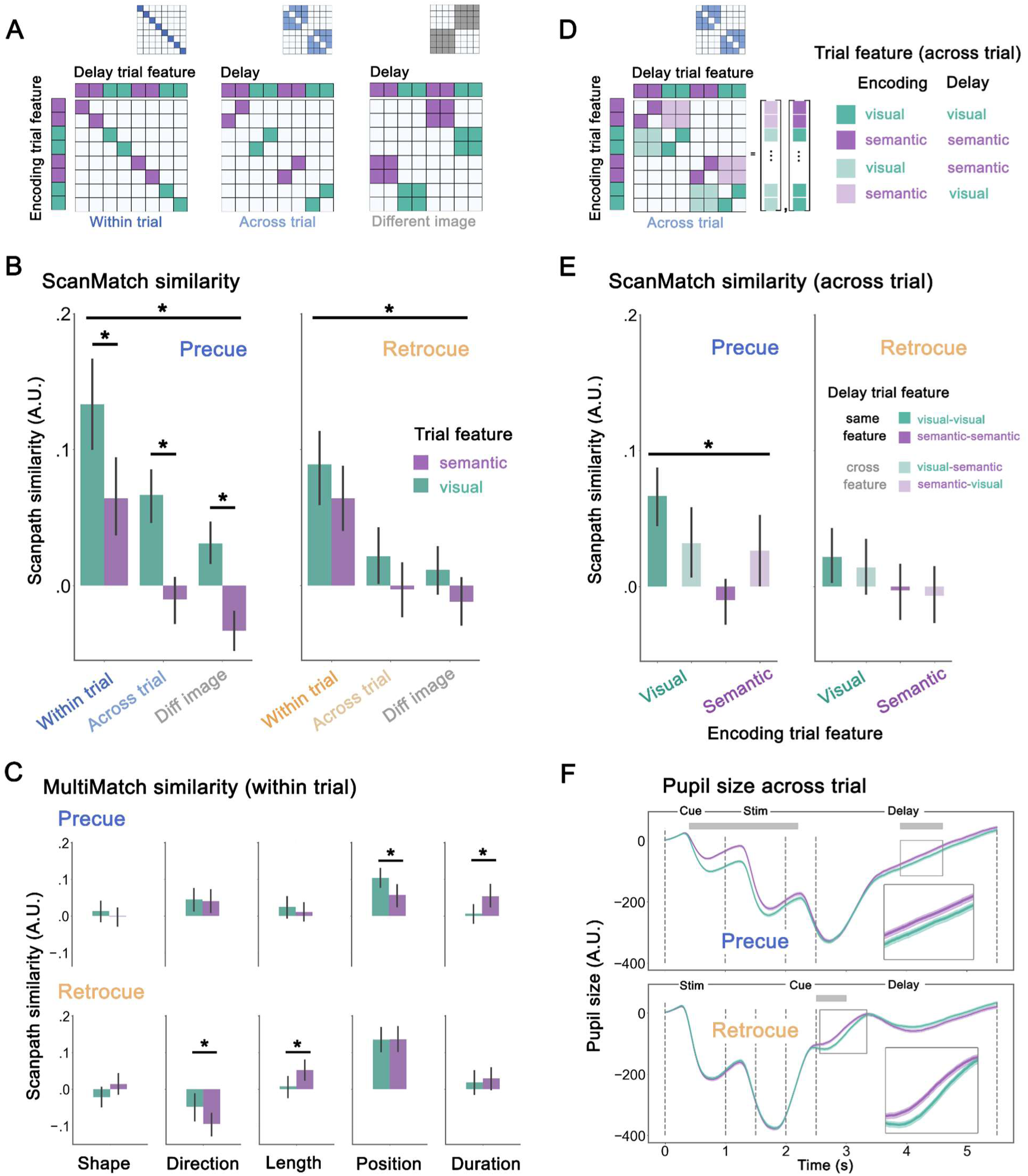
Scanpath similarity between encoding and delay epochs, across visual and semantic trial conditions. (**A**) A similar matrix schematic as in Figure 2, now illustrating visual (green) and semantic (purple) conditions, for two different images. The first 4 rows/columns all reflect trials with the same sample image, while the second half reflect a different sample image. Scanpath similarity scores for each comparison condition (“within trial”, “across trial”, “different image”) were partitioned into visual or semantic trial conditions. (**B**) The mean of individual subject scanpath similarity scores (ScanMatch algorithm) across comparison conditions and trial types, in precue and retrocue experiments. Note the similarity matrices were normalized within subject to account for baseline variability. Horizontal lines across the top of the plot indicate a main effect of comparison condition, whereas shorter lines accompany significant post-hoc pairwise comparisons. Note that the precue experiment also showed both a main effect of trial condition as well as an interaction between factors. (**C**) Scanpath similarity scores for visual and semantic trials, across five geometric dimensions (MultiMatch algorithm), for “within trial” scanpath comparison. (**D**) Schematic for cross-feature comparisons within the “across trial” condition. Darker shades represent congruent encoding and delay trial types (visual-visual, semantic-semantic), and lighter shades represent incongruent (visual-semantic, semantic-visual) (**E**) Encoding-delay scanpath similarity scores across trials of either the same type (visual or semantic), or across trial types. Note that the “same feature” condition depicts the same data as in the “across trial” condition of panel B, but here it is contrasted with scanpath similarity between trials that emphasized different features. Significance bar represents an interaction between encoding trial condition and delay trial congruence. (**F**) The timecourse of average pupil size across the trial, for visual and semantic trial types, in precue and retrocue experiments. Grey bands at top of plots indicate periods where permutation testing yielded a significant difference between visual (green) and semantic (semantic) conditions.

Lastly, we zeroed in on the “across trial” condition to examine scanpath similarity between the *visual* and *semantic* trial types. The idea here was to examine whether generalizable image-specific information in gaze patterns would only be present across trials of the same type (i.e., only within visual trials), or whether one trial type might encourage an encoding strategy that better generalizes between trial types (e.g., *visual* encoding may share scanpath similarity with a *semantic* delay, but not the other way around). Therefore, in addition to selecting encoding-delay pairs within the same trial feature (as in the preceding analyses), here we also selected pairs where the encoding and delay came from different trial types. This generated four conditions, crossing each encoding and delay trial type: *visual-visual, visual-semantic, semantic-visual, or semantic-semantic* (**Figure 3D**). As with the other scanpath analyses, similarity values within each condition were averaged to produce one value per condition, per trial.

### Representational similarity analysis

In the final analysis, we assessed the timecourse of stimulus representation in gaze patterns through a Representational Similarity Analysis (RSA). RSA is often applied to multi-voxel fMRI activity patterns to characterize the stimulus information carried in physiological signals (Kriegeskorte et al., 2008). RSA is geared to gauge the representational information in lower-dimensional structure, even when specific activity patterns vary. For instance, two similar stimuli might be expected to show activity patterns that remain similarly correlated over time, even if the underlying activity for both stimuli changes. This approach is therefore well-suited to extract stable representational information over the course of a WM task, when activity may dynamically fluctuate. This approach has also been applied to patterns of microsaccade bias during WM, to show that they reflect the relational structure between stimuli of different orientations (Linde-Domingo & Spitzer, 2024).

However, the similarity between our complex naturalistic stimuli is less straightforward to quantify. While a single feature like orientation varies along just one dimension, natural scenes can vary along myriad dimensions. We therefore used VGG-16, a convolutional neural network specialized for image recognition, to glean the similarity structure of our stimulus space (Simonyan & Zisserman, 2015). We computed pairwise Pearson correlations for every image pair, based on their activity in each layer of the model (similar to Piriyajitakonkij et al., 2024). Specifically, the modeled similarity for a layer was calculated as

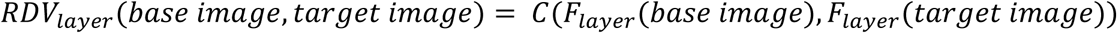

Where *F_layer_* is the activity in a given layer of the model in response to an image, and C is the Pearson product-moment correlation coefficient. These pairwise correlations produced a model-derived similarity matrix (**Figure 4A**), which we compare to each participant’s empirically-derived gaze similarity matrix.

**Figure 4.**
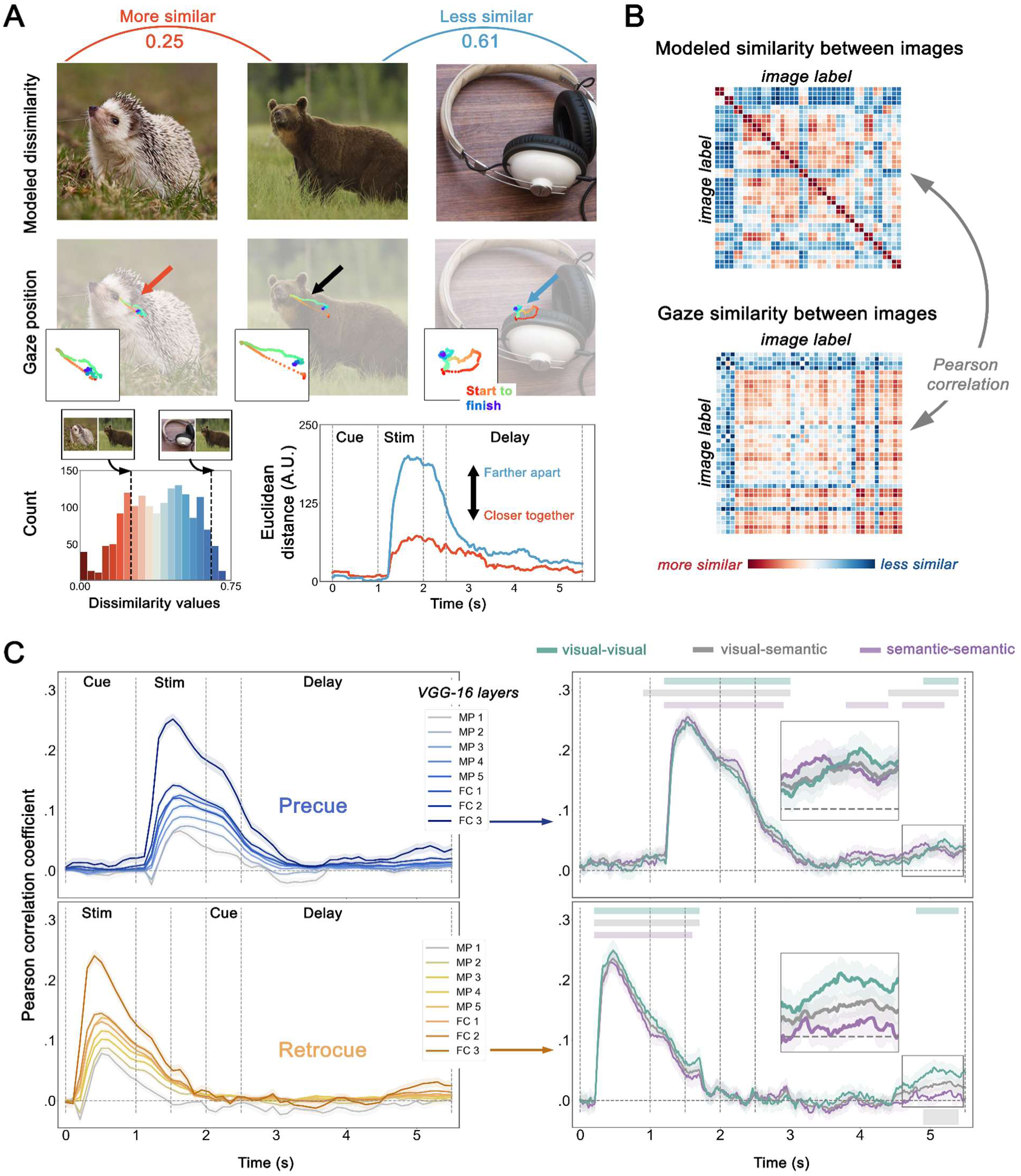
Representational similarity analysis methods and results. (**A**) Evaluating stimulus similarity and gaze similarity for representational similarity analyses (RSA). Top panel: example similarity values between image pairs, as assessed via correlation in the final layer of VGG-16. The first pair [bear, hedgehog] was rated as relatively more similar, and the second [bear, earphone] as less similar, according to their activity patterns in the network. Below that are example gaze coordinates during WM for the corresponding image. Cooler colors represent later time points. Lower panel: The distribution of scores, corresponding to the similarities between all unique image pairs (left), and an example time course of Euclidean distances in gaze between those image pairs. (**B**) Top: matrix depicting pairwise similarity scores for every image pair in the stimulus set, assessed via correlations between VGG-16 layer activity for each image. Bottom: average gaze similarity (Euclidean distance) for respective image pairs, collapsed across time. (**C**) Pearson correlation between gaze similarity and modeled similarity, across every trial time point, evaluated for key max pooling (MC) and fully connected (FC) VGG-16 layers. Darker lines represent later layers of the model. (**D**) Pearson correlation between gaze similarity and modeled similarity (from activity in layer FC3), split by trial comparison conditions (visual-visual, visual-semantic, and semantic-semantic). Thin shaded bands on top of plot show periods when correlation values were greater than chance, for each trial condition. Thicker band at bottom of plot shows periods when trial conditions differed from each other.

To construct the gaze similarity matrix **(Figure 4B**), we used only the 5.5s period of interest (i.e. fixation offset to the end of memory delay), then downsampled to 10Hz for efficiency. For every participant, we calculated similarity as the Euclidean distance in gaze coordinates between every trial pair, for every time point (cf. Linde-Domingo & Spitzer, 2024). Then we averaged across any pairings of the same two images.

Because there were 40 unique stimuli, this produced one 40 x 40 model-derived image similarity structure, and a 40 x 40 gaze similarity structure for each of 55 timepoints. Note, the raw values of the modeled similarity ranged from −1 to 1, with larger values signifying maximum similarity (same image). In contrast, in gaze, smaller values in Euclidean distance indicated higher similarity. To align the direction of modeled stimulus similarities and gaze pattern, we normalized the model output to the interval [0,1], then subtracted that value from 1, so that 0 represented the strongest similarity (same image). We then computed the Pearson correlations between the image and gaze similarity structures for each timepoint (**Figure 4B**). This produced one time series of Pearson correlation for each participant, indexing the degree to which their gaze patterns followed the similarity structure of the stimulus space. When the two structures are correlated above chance, we can say that the gaze pattern conveys stimulus-specific information.

We first examined the stimulus information in gaze as indexed across several layers of the neural network model. We conducted the RSA based on activity in the five max pooling layers and three fully connected layers of VGG-16 (**Figure 4C**). Since the modeled similarity from the last layer (FC3) exhibited the highest correlation with the gaze structure, we used only this last layer for subsequent analyses.

We then created separate gaze similarity structures for three conditions: correlations between *visual-to-visual*, *semantic-to-semantic*, or *semantic-to-visual* trials. Note that here, unlike the scanpath analysis, the *semantic-visual* and *visual-semantic* conditions are identical. We then averaged across any pairings of the same images within each condition, creating three 40 x 40 x 55 matrices. Then we carried out the same RSA correlation (with the modeled image similarity) separately for each comparison condition.

We conducted permutation testing on the RSA time series, similar to previous work (Dong & Kiyonaga, 2024; Zokaei et al., 2019). To test whether correlations for each condition were significantly above chance, we first randomly flipped the signs of half of the participants’ correlations in each iteration. Then we tested the permuted correlations against zero via a one-sample *t-*test, once for each time point. From the resulting array of p-values, we found the largest cluster that exceeded the significance threshold (*p* < 0.05) and summed the t-values corresponding to this cluster. Then, this procedure was repeated 5000 times to generate a permutation distribution for each condition. The same *t-*test was run using the actual correlation scores, and the clusters of significance were identified. The *p*-value of each cluster could be calculated by comparing the actual tsum of the true clusters versus the permutation distribution as P(T > *t*sum). If this value is smaller than 0.05, then such a cluster in the correlation trace would be considered significantly above zero.

To test if the gaze RSA correlation differed between the trial-type comparison conditions, we ran a cluster-based permutation ANOVA, where we first shuffled the condition labels (*visual-visual, semantic-semantic,* and *semantic-visual*). Then we applied a one-way ANOVA to each time point, with the shuffled label as the independent variable, and correlation value as the dependent variable. Instead of t-values, here we recorded the sum of f-values of the largest cluster that exceeded significance. This procedure was likewise repeated 5000 times to generate the permutation distribution. We calculated the *F* and *p* values of the real (not permuted) data and identified clusters that exceeded significance. Then the Fsum of these clusters was compared to the permutation distribution.

### Pupil analysis

For pupil data, artifact rejection followed the methods described in the *Preprocessing* section, and we then applied the same interpolation procedure for sections of missing data. While we used linear or nearest neighbor interpolation to reconstruct the gaze data, we used linear and cubic spline for pupils. The slightly different interpolation methods for the two metrics is meant to better capture their respective physiology (Mathôt & Vilotijević, 2023). Then we also removed any trials that had been excluded based on gaze data quality.

To compare the pupil size under visual and semantic task conditions, we first subtracted each trial timeseries from a baseline value. The baseline was taken as the average pupil size during the last 200ms of the fixation period at the beginning of each trial. We then partitioned the data based into *visual* vs. *semantic* trials.

We applied cluster-based permutation *t-*tests to identify periods of significant difference in pupil size between *visual* vs. *semantic* trials. In each iteration, we randomly shuffled the trial condition labels, and applied independent *t*-tests at each timepoint using the shuffled label as the independent variable. The rest of the procedures followed the steps outlined in the previous sections.

## Results

### Working memory recognition performance

WM recognition performance was overall good (mean accuracy: 90.49% correct), and did not differ between pre- and retrocue experiments (*accuracy*: *t*(80) = 1.64, *p* = 0.11, *Cohen’s d* = 0.36; *RT*: *t*(80) = −0.48, *p* = 0.63, *Cohen’s d* = −0.11). However, performance was somewhat better on *semantic* trials (vs. *visual*; **Figure 1D**). In both experiments, when the WM category was probed instead of the visual detail, responses were more accurate (precue: *t*(40) = −8.78, *p* < 0.001, *Cohen’s d* = −1.38; retrocue: *t*(40) = −5.69, *p* < 0.001, *Cohen’s d =* −0.86) and faster (precue: *t*(40) = 4.78, *p* < 0.001, *Cohen’s d* = 0.27; retrocue: *t*(40) = 4.89, *p* < 0.001, *Cohen’s d* = 0.36). Consistent with previous findings, therefore, we observed a semantic task advantage (Brown & and Wesley, 2013; Lee et al., 2013), but the two different experiments were comparable to each other.

### Gaze follows an image-specific scanpath during working memory delays

Our first goal was to assess whether the gaze patterns during a WM delay recapitulated the encoding patterns for the same image. **Figure 1C** illustrates some example gaze heatmaps that vary in similarity to each other from encoding to delay. However, there are many ways to quantify these patterns. To capture both the spatial and temporal properties of the gaze signals, here we first examined whether scanpaths — sequences of successive eye movements — were aligned between encoding and delay epochs (**Figure 2A** and **B**).

Our first analyses use the string-based approach, assessing similarity between two scanpaths via the **ScanMatch** algorithm (Coutrot et al., 2018). We compared the strength of encoding-delay similarity for three scanpath comparison conditions: within the same trial (‘within trial’), between different trials when the same image was remembered (‘across trial’), and between different trials when a different image was remembered (‘different image’; **Figure 2B** and **C**). If gaze patterns during WM convey image-specific spatio-temporal information, then encoding-delay scanpath similarity should be greater across trials when the same image is remembered, as compared to trials when different images are remembered.

We ran a one-way repeated measures ANOVA on ScanMatch similarity scores, with a 3-level factor of scanpath comparison condition: “within trial”; “across trial”; “different image”. In both experiments, we found that scanpaths during the WM delay resembled encoding scanpaths for the same image (**Figure 2D**). Encoding and delay scanpaths were most similar to each other within the same trial, then across trials with the same sample image, followed by different trials with different sample images (precue: *F*(2,80) = 46.15, *p* < 0.001, ηp2 *=* 0.54; retrocue: *F*(2,80) = 92.75, *p* < 0.001, ηp2 *=* 0.70). Follow-up *t*-tests confirmed that ‘within trial’ similarity scores were greater than ‘across trial’ (precue: *t*(40) = 5.49, *p* < 0.001, *Cohen’s d* = 1.17; retrocue: *t*(40) = 8.95, *p* < 0.001, *Cohen’s d* = 1.93), but ‘across trial’ scores were also greater than ‘different image’ scores (precue: *t*(40) = 9.82, *p* < 0.001, *Cohen’s d* = 2.22; retrocue: *t*(40) = 2.92, *p* = 0.006, *Cohen’s d* = 0.59). Thus, shared gaze patterns during WM are not limited to a particular trial, but can carry image-specific information that generalizes over different trials, if the same image is being remembered.

Gaze patterns during a blank WM maintenance period may (at least partly) retrace the unique, image-specific paths that the eyes took during encoding. We will refer to this encoding-delay similarity as ‘WM gaze reinstatement’, akin to the shared encoding-retrieval gaze patterns that have been observed during long-term memory (Johansson & Johansson, 2014; Wynn et al., 2019).

### WM gaze reinstatement is stronger when the task goal is known before encoding

Here, participants were cued as to the task-relevant feature either before or after encoding the WM sample image (i.e., precue vs. retrocue). We reasoned that this experimental difference might also alter encoding-delay gaze similarities, if participants adopt different strategies in the different tasks. For instance, participants might take a more focused encoding approach in the precue experiment, to extract only the most relevant information. However, they may need to encode more dimensions of information in the retrocue experiment, because the probe demands are not yet known. This could yield stronger encoding-delay gaze similarity in the precue experiment.

Indeed, encoding-delay scanpath similarity scores appeared overall lower during the retrocue task (**Figure 2D**). To investigate this difference, and ensure that eye movements weren’t overall lower during the retrocue task, we examined additional gaze metrics. We first examined saccade frequencies, across the whole trial epoch, in both experiments (**Figure 2E**). Saccade frequencies were descriptively elevated overall in the retrocue task (vs. precue), but especially during the WM sample encoding period. Saccade frequency was higher during encoding (as compared to delay) in only the retrocue task (precue: *t*(40) = 0.75, *p* = 0.46, *Cohen’s d* = 0.11; retrocue: *t*(40) = 2.56, *p* = 0.01, *Cohen’s d* = 0.38). Rather than a reduction in eye movements that could explain dampened encoding-delay similarity in the retrocue experiment, participants may take a different approach to encoding the WM sample.

To examine this further, we also tested for ‘WM gaze reinstatement’ using vector-based metrics that quantify multiple distinct scanpath properties. The **MultiMatch** algorithm (Dewhurst et al., 2012; Jarodzka et al., 2010; Wagner et al., 2019), evaluates eye movement sequences along the dimensions of shape, direction, length, position, and duration (**Figure 2F**).

In the precue experiment, all metrics except for shape exhibited WM gaze reinstatement within trials (**Figure 2F**; See **Supplementary Table 1** for statistics). However, the image-specific gaze reinstatement across trials was only present in the position metric (see **Supplementary Table 2** for statistics). This indicates that gaze tended to visit the same coordinate positions whenever the same image was remembered. The retrocue experiment similarly showed within-trial gaze reinstatement for all metrics except shape, and between trials for only the position metric. However, the direction metric showed that scanpaths were inversely related to each other from encoding to delay. That is, rather than the positive relationship that was observed between encoding-delay scanpaths in the precue experiment, the scanpaths showed a negative relationship in the retrocue experiment. Despite gaze reinstatement in position during the delay, the angular direction of the saccade vectors were more like the opposite of those during encoding. This could arise if gaze revisited the same coordinate positions as during encoding, but in a different order.

In sum, the timing of task cues may influence WM encoding strategies, and the nature of encoding-delay reinstatement in gaze. Saccades were overall more frequent during retrocue task encoding, but gaze position showed image-specific encoding-delay similarity in both experiments. However, the direction gaze metric showed sharply different patterns between the pre- and retrocue experiments, suggesting that eye movements may have followed a reversed temporal sequence when the task condition was cued after encoding.

### WM gaze reinstatement is stronger when the task requires visual detail

A primary goal of this study was to test whether WM gaze patterns would be flexible to the expected demands of an upcoming WM test, even when the remembered stimuli were physically identical. We therefore examined whether WM gaze reinstatement – as indexed by encoding-delay scanpath similarity – differed when the task emphasized visual vs. semantic maintenance strategies.

We first assessed encoding-delay similarity within and between trials (as in **Figure 2D**), but separately for *visual* and *semantic* trial conditions (**Figure 3A**). For both experiments, we ran a repeated-measure ANOVA on ScanMatch similarity scores with factors of scanpath comparison (“within trial”, “across trial”, “different image”) and trial condition (*visual* vs. *semantic*). Like earlier analyses, both experiments showed a main effect of scanpath comparison, whereby encoding-delay similarity was greatest within trial, then across trials with the same image, then across trials with different images (precue: *F*(2,80) = 45.45, *p* < 0.001, ηp2 = 0.53; retrocue: *F*(2,80) = 91.10, *p* < 0.001, ηp2 *=* 0.70). Additionally, the precue experiment showed an effect of trial condition (*F*(1,40) = 13.72, *p* = 0.001, ηp2 *=* 0.26), and an interaction between factors (*F*(2,80) = 4.03, *p* = 0.02, ηp2 *=* 0.01; **Figure 3B)**. Encoding-delay scanpath similarity was overall stronger for *visual* trials (vs. *semantic*) and the difference between visual and semantic strategies was most pronounced within trials. Therefore, WM gaze reinstatement was relatively amplified when the WM probe encouraged a precise visual memory strategy. However, in the retrocue experiment, there was no effect of trial condition, nor an interaction between factors (trial condition: *F*(1,40) = 0.46, *p* = 0.50, ηp2 *=* 0.01; interaction: *F*(2,80) = 1.76, *p* = 0.18, ηp2 *=* 0.04). Therefore, the gaze reinstatement difference between *visual* and *semantic* trials only emerged when the cue allowed a task-specific strategy during encoding.

The vector-based MultiMatch results also followed this general pattern and the pattern from earlier analyses (**Figure 3C**; **Supplementary Figure 1** shows all scanpath comparison conditions). The difference between *visual* and *semantic* trial gaze reinstatement was evident only in the position metric for the precue task (interaction: *F*(2,80) = 3.81, *p* = 0.03, ηp2 *=* 0.09), and the retrocue task showed an inverse pattern in the direction metric from encoding to delay (interaction: *F*(2,80) = 3.68, *p* = 0.03, ηp2 *=* 0.08; **Supplementary Table 3** and **4** show all conditions and statistics). The difference between *visual* and *semantic* trials was not explained by differences in overall rates of eye movement, as saccade frequencies did not differ between trial conditions (**Supplementary Figure 2**). Therefore, the extent and type of WM gaze reinstatement was flexible to both task- and trial-level strategy.

The effect of trial condition on encoding-delay scanpath similarity (i.e., **Figure 3B**), suggests that scanpaths only show generalizable image-specific information when the task encourages a visual strategy. However, this effect could arise because a visual encoding strategy is always more generalizable to other trials, or instead because only visual trials show strong alignment from encoding to delay. Therefore, we also tested encoding-delay scanpath similarity across the two trial types – for instance, pairing a visual encoding trial with a semantic delay trial (**Figure 3D**). We did this for only the ‘same image, across trial’ comparisons, to gauge which conditions yielded the most generalizable image-specific gaze patterns across trials (**Figure 3E**; See **Supplementary Figure 3** for “different image” comparisons). In the precue task only, we found that any combination of trials that included the *visual* strategy (as either the encoding or delay input) produced superior scanpath similarity scores (**Supplementary Table 5**). Scanpath similarity was strongest when comparing encoding from *visual* trials to delay from *visual* trials, but encoding-delay comparisons across trial features (*visual-semantic* or *semantic-visual*) were stronger than comparisons between only *semantic* trials. Therefore, image-specific information in gaze patterns is minimal when the task encourages a semantic strategy.

The timecourse of pupil dilation also corroborated task- and trial-level strategy modulations (**Figure 3F**). Pupils can be expected to constrict at the onset of a new visual stimulus (Mathôt et al., 2018; Oster et al., 2022), and during fine-grained visual discrimination (Ebitz & Moore, 2019; Einhäuser, 2017). Here, we found that stimulus-induced constriction was also modulated by task context. In the precue task, the constriction during encoding was greater on *visual* trials, and pupil size also remained smaller on *visual* trials throughout the WM delay. In the retrocue task, pupil size was identical throughout encoding (as expected when the trial feature is not yet known), but *visual* and *semantic* trials diverged shortly after the retrocue offset. Even in the absence of a visual stimulus, WM maintenance (precue) and internal attentional selection (retrocue) showed relative pupil constriction, consistent with visual discrimination, when the upcoming probe would require more fine detail.

Together, gaze and pupil metrics indicate that the same WM content may be subject to different encoding and maintenance strategies when the task context and probe demands differ. WM delay period gaze patterns held closer to encoding patterns when visual details were emphasized – but only when the task strategy was cued before encoding. Image-specific eye movement patterns, during a blank WM delay, were modulated by upcoming WM task demands.

### Gaze tracks the similarity relationships between naturalistic images during WM

The prior analyses have shown that gaze patterns during the WM delay share properties with the gaze patterns made during stimulus encoding, and that this similarity is accentuated when the task encourages a more visual encoding approach. However, even in cases when eye movement patterns differ from encoding to delay, they may still convey stimulus-specific information. For instance, when stimulus locations change from encoding to probe, location-specific WM gaze biases can shift to prospectively reflect expected future locations (Boettcher et al., 2021; Liu et al., 2024). To examine the timecourse of stimulus representation in the gaze patterns – and when visual vs. semantic differences might arise – these final analyses take a representational similarity analysis (RSA) approach.

We constructed matrices for both model-derived image similarity and empirically-derived gaze similarity (**Figure 4A)**. When the two structures are correlated (**Figure 4B**), we can say that the gaze pattern conveys stimulus-specific information. We found that, for both pre- and retrocue tasks, image similarity from the last layer of the model showed the strongest correlation with the gaze data (**Figure 4C**). Such later layers reflect the most high-level image information, suggesting that WM gaze patterns similarly express the most integrated visual representations (rather than particular low-level components). The remaining analyses use the similarity structure from that final layer.

We created gaze similarity structures for comparisons between *visual-to-visual*, *semantic-to-semantic*, or *semantic-to-visual* trial pairs. We separately ran the RSA for these three comparisons and found that representational information was strongly present during encoding in all conditions, in both the pre- and retrocue experiments (**Figure 4C**). In other words, the correlations between trials in the gaze structure resembled the model-derived similarity between stimuli. More similar WM sample stimuli showed more similar gaze patterns. Permutation testing confirmed that this correlation was above chance during encoding in all three comparison condition (*visual-visual*: 1.2s to 3s, *p* < 0.001; *semantic-semantic*: 1.2 s to 2.9 s, *p* < 0.001; *semantic-visual*: 0.9s to 3s, *p* < 0.001). Image-specific gaze patterns during WM encoding therefore reflected the relational structure between these naturalistic images.

Stimulus information tapered off after encoding, but ramped back up again toward the end of the delay. In both experiments, WM stimulus information in the gaze pattern – at least as captured by the VGG-16 similarity structure – emerged only late in the WM delay, as the probe approached. In the precue experiment, representational information was present late in the delay for all three conditions (*visual-visual*: 4.9s to 5.4s, *p =* 0.04; *semantic-semantic*: 3.8s to 4.4s, *p* = 0.03; 4.6s to 5.2s, *p* = 0.03; *semantic-visual*: 4.4s to 5.4s, *p* = 0.02). However, in the retrocue experiment, this late-delay escalation occurred only in the *visual-visual* condition (4.8s to 5.4s, *p* = 0.02). No significant clusters were detected in the *semantic-semantic* condition, although one cluster approached significance in the *semantic-visual* condition (5.5 s to 5.9 s, *p* = 0.06). An additional permutation analysis tested the difference between trial feature comparisons (*visual-visual, semantic-semantic, semantic-visual*) and found an effect in only the retrocue experiment (5.4 s to 5.9 s *p* = 0.04). Stimulus-specific information emerged late in the retrocue WM delay for only *visual* trials.

RSA further showed that gaze patterns reflect image-specific information during WM, even for complex naturalistic images. At various points across a WM trial, stimuli that share more similar image properties also evoke more similar gaze patterns. The high-level image properties that determine similarity in a deep convolutional neural network also shape gaze patterns during WM.

## Discussion

Across two experiments we asked participants to remember real-world images, and we manipulated whether the visual details or semantic labels were most relevant for the WM test. We assessed WM gaze reinstatement, operationalized as the similarity in spatio-temporal eye movement sequences (scanpaths) from stimulus encoding to WM delay. We also used deep convolutional neural network modeling to guide RSA of the evolving stimulus information in gaze patterns across a trial. Across analyses, we found that eye movement patterns during WM conveyed image-specific information that generalized across trials. This WM gaze reinstatement was most pronounced when the task required greater visual detail, and when the task strategy was cued before stimulus encoding. Together, the findings reveal precise, but context-sensitive, expressions of mnemonic content in oculomotor inflections.

### Distributed working memory representations across the nervous system

The anatomical substrates and representational nature of WM content have been the subject of ongoing debate (Leavitt et al., 2017; Sreenivasan et al., 2014; Xu, 2017). Much human neuroimaging supports the idea that WM maintenance engages overlapping neural representations with sensory perception (D’Esposito & Postle, 2015; Serences et al., 2009), but it has become increasingly clear that WM-related information is widely distributed across the brain (Christophel et al., 2017; Courtney et al., 1997; Dotson et al., 2018). Beyond the cortex, signatures of basic WM content properties may also be reflected in the peripheral sensorimotor apparatus (Dong et al., 2025; Dong & Kiyonaga, 2024; Hustá et al., 2019; Kay et al., 2022; van Ede et al., 2019; Zokaei et al., 2019; Yang et al., 2025). Here, we find that feature-precise and flexible WM content information is present in the eyes. More than simple location tagging, gaze patterns during WM retrace spatio-temporal sequences associated with specific naturalistic images. This WM gaze reinstatement is more than a lingering sensory-evoked trace, or reflexive response to visual stimulation, as it was acutely sensitive to which stimulus dimensions were needed for the task. WM may adaptively recruit the relevant sensory and motor structures across the nervous system to support current needs (Postle et al., 2006).

### Working memory encoding and delay strategies vary with context

A hallmark of WM is its flexibility. Humans can strategically shift and spread attention between the inner and outer worlds or among items held in mind, and can flexibly use mental content toward a variety of goals (Gazzaley & Nobre, 2012; Nobre & Stokes, 2019; Souza & Oberauer, 2016). However, the representations that support such flexible use have been contested (Oberauer & Awh, 2022; Stokes, 2015). Here, we varied contextual WM demands at the level of the overarching task (precue vs. retrocue) and at the trial level within tasks (*visual* vs. *semantic*), to examine how oculomotor WM adapts to these situational factors.

We found that both the stimulus encoding and maintenance strategies (indexed by gaze patterns) differed depending on when the task-relevant features were cued. In a dynamic and uncertain visual environment, exploration behavior would be most beneficial (Cohen et al., 2007). In our case, when the relevant task features were unknown during stimulus presentation (i.e., retrocue task), the task may favor a more holistic, exploratory sampling strategy. This fits with the relatively increased saccade frequency that we observed during retrocue task encoding. During the precue task, on the other hand, participants know before encoding which stimulus feature they should prioritize. When encoding can be proactively targeted to a particular task strategy, WM maintenance may sustain that strategy, yielding more similar gaze patterns from encoding to delay.

In addition to different encoding strategies, therefore, the task context might determine what kind of processing occurs during maintenance. In the retrocue task, the encoded information may undergo a representational transformation in the delay (Souza & Oberauer, 2016). We found that, when the relevant WM content had to be retroactively selected from memory, eye movement patterns were anti-correlated from encoding to delay in some gaze metrics. This may echo the reversed direction of information flow that is observed from LTM encoding to retrieval (Linde-Domingo et al., 2019).

Together, these findings suggest that oculomotor WM signatures reflect flexibility in encoding-delay strategies. When the task mode is known before encoding, the eyes may sample information selectively and maintain a stable representation during the delay (especially for visual detail). When the task-relevant features are unknown at encoding, the eyes may sample more holistically but then incur a task-oriented representational shift during the delay.

### Ocular working memory signatures anticipate task-relevant features

In addition to the strategic differences provoked by cue timing, oculomotor WM signatures also varied with the particular stimulus dimension that was cued. When the WM test required more precise visual memory, delay period gaze resembled the encoding pattern more strongly. Given the same encoded stimulus, the delay-spanning eye movement patterns depended on the expected upcoming task demands. This aligns with neuroimaging findings that visual cortical regions are preferentially engaged during WM for more visual (vs. semantic) features (Lee et al., 2013). Likewise, we speculate that our observed difference in WM gaze reinstatement may reflect a modal difference in the circuitry engaged between *visual* and *semantic* conditions. We found that *visual* trial scanpaths seemed to generalize across trial types better than *semantic* trial scanpaths, which might occur if gaze patterns under a visual strategy contain more image-specific signal, in general. Moreover, stimulus information in the RSA ramped up in anticipation of the probe, especially in the visual condition. This further corroborates the idea that oculomotor engagement during WM is prospective and task-dependent (Ball et al., 2013; McAteer et al., 2023; van Ede et al., 2021), rather than retrospective content storage.

We also found that pupil size was relatively smaller during the WM delay when the task encouraged more detailed visual memory. Smaller pupil size has been shown to benefit fine-grained discrimination, while larger pupil size benefits detection (Eberhardt et al., 2022; Ebitz & Moore, 2019; Mathôt et al., 2018; Mathôt & Ivanov, 2019). Our findings suggest that such pupillary mechanisms for perception are also reflected during ‘internal’ visual discrimination. Together, these results show that various ocular inflections during WM are modulated by the task-relevant stimulus features. Humans may remember the same information differently depending on what they need to do with it (cf. Trentin et al, 2023, 2024; Henderson et al., 2022; Nobre & Stokes, 2019).

### Multiple routes to successful working memory

It is unclear whether eye movements during WM play any causal role in maintenance. There is some evidence that interfering with eye movements can interfere with WM (Postle et al., 2006) and LTM (Johansson & Johansson, 2014, 2020), but the link is tenuous (Liu et al., 2022; Loaiza & Souza, 2022; Walcher et al., 2024). Here, we can say that the eye signals are functionally modulated, in that they vary depending on the task-relevant features. But pre- and retrocue task performance metrics were matched while their associated oculomotor signatures were distinct, which might suggest that the eye movements are not strictly essential. Instead, there can be multiple pathways to the same performance outcome. Indeed, people with different preferred maintenance strategies (e.g., visual vs. verbal) may vary in their ocular engagement (Korda et al., 2024) but be equally capable of most WM tasks (Keogh & Pearson, 2018; Korda et al., 2024).

The current findings have both practical and theoretical implications. This work suggests that gaze may be a feasible tool to decode the properties of what is being remembered, or to gauge the integrity of different underlying representations (e.g., sensory vs. linguistic). The findings also suggest that the representations underlying WM may differ depending on the resolution required for the task at hand. Therefore, our understanding of visual WM storage based on lab tasks that test for precise visual features may not apply to ‘good enough’ WM that can be achieved with a semantic label or category code. Indeed, performance was overall better in the *semantic* condition here, raising concerns that the findings could stem from greater task difficulty in the *visual* condition. If that were the case, however, we would have expected relatively greater pupil dilation (rather than smaller) during the *visual* trials (Kahneman & Beatty, 1966; van der Wel & van Steenbergen, 2018). Nonetheless, future work should seek to determine whether there is an obligatory link between WM and eye movements, when difficulty is controlled.

### Concluding remarks

Information that we hold in WM can be detected from the eyes – an outward extension of the brain and the first stop for visual perception. We found that, during WM maintenance, eye movements recapitulate the spatiotemporal sequences made during stimulus encoding. This WM gaze reinstatement carries not only basic information about item location, but also image-specific patterns for complex natural scenes. Most critically, we see that the WM oculomotor signal is not merely a reflexive sensory reinstatement, but also varies with WM strategy and prospective task demands. In this sense, the eyes make an informative, accessible readout of otherwise hidden mental representations. WM is not confined to cortical representation, but flexibly extends out to the earliest stages of sensorimotor processing.

## Acknowledgements

We thank research assistants Connie Xie for help with data collection, as well as Ana Chkhaidze, Pria Daniel, Sihan Yang, and Zhuojun Ying for helpful comments on the manuscript. This work was partially supported by NIH award 1R01EY036843-01 to A.K as well as the Air Force Office of Scientific Research (AFOSR) under award number FA9550-22-1-0230.

## Data and code availability

Study materials and provisional code are available here: https://github.com/YueyingDong/dong_etal_paper_code/tree/main. Provisional datasets are available here: https://osf.io/PYEBZ/.

## Notes

### Competing Interest Statement

The authors have declared no competing interest.

### Summary of Updates

Revised Abstract and Introduction to reflct a more recent submission version.

